# A Quantitative Model of Cellular Decision Making in Direct Neuronal Reprogramming

**DOI:** 10.1101/2020.02.04.933713

**Authors:** Adriaan Merlevede, Viktor Drugge, Roger A. Barker, Janelle Drouin-Ouellet, Victor Olariu

## Abstract

The direct reprogramming of adult skin fibroblasts to neurons is thought to be controlled by a small set of interacting gene regulators. Here, we investigate how the interaction dynamics between these regulating factors coordinate cellular decision making in direct neuronal reprogramming. We put forward a quantitative model of the governing gene regulatory system, supported by measurements of mRNA expression level dynamics. We find that reinterpreting the interaction between two genes (PTB and nPTB) is necessary to capture quantitative gene interaction dynamics and correctly predict the outcome of various overexpression and knockdown experiments. This analysis is strengthened by a novel analytical technique to dissect system behaviour and reveal the influence of individual factors on resulting gene expression. Overall, we demonstrate that computational analysis is a powerful tool for understanding the mechanisms of direct (neuronal) reprogramming, paving the way for future models that can help improve cell conversion strategies.

## Introduction

Cell differentiation, the process that establishes cellular identity, traditionally follows a well-established hierarchy from totipotent to nilpotent cell types. However, the past decade has seen great advances in reprogramming procedures, allowing researchers to convert terminally differentiated cells to other cell types *in vitro*. This technology has promising applications for disease modelling, where an individual’s skin fibroblasts can be converted to neurons and be used to determine their own unique neuronal pathology *in vitro*, as well as regenerative medicine, where replacement tissues can be sourced from a patient’s own body. Direct reprogramming, in particular, allows conversion between terminally differentiated cell types without passing through an intermediate pluripotent state. This gives the opportunity to obtain mature neurons quickly (within a month), allowing for relatively easy handling of dozens of lines by one experimenter. Moreover, they are not clonal, thus avoiding the risk of clonal bias. Importantly, multiple reports have recently demonstrated that directly reprogrammed neurons retain many important aspects of the age signatures of the donor, including age-related changes in the epigenetic clock, the transcriptome and microRNAs, the reactive oxygen species (ROS) levels, DNA damage and telomere lengths, as well as the metabolic profile and mitochondrial defects^1,2,3,4^. As these cellular changes are suspected to play crucial roles in the development of age-associated disorders, this makes direct neuronal reprogramming an ideal approach to model neurodegenerative diseases for which age is the most important risk factor. However, because the source cells do not pass through a proliferating intermediate stage, a high conversion efficiency is required for large scale clinical application.

The working principle of contemporary reprogramming methods is to introduce a select combination of transcription factors and other biomolecules that manipulate the expression of key genes, initiating a cascade of regulatory mechanisms that control all known aspects of cell identity. Finding a combination of appropriate factors that induce the desired conversion with sufficient efficiency is the central difficulty for current reprogramming research, and is typically done by trial and error, using a set of candidate genes picked from databases and published resources based on the characteristic expression profiles of the source and target cell types. Some effort has gone into streamlining this process of selecting candidate genes^5^, but the efficiency of these tools is bounded by our knowledge of the underlying gene interactions.

One of the most notable cases is conversion of somatic dermal cells into neurons and shows great promise in the study and treatment of neurodegenerative disorders^6^. Dermal fibroblasts are highly suitable source cells because they are abundant and easy to harvest. Many different cocktails of transcription factors have been successfully applied. Originally, Brn2, Ascl1 and Myt1l were shown to be sufficient in adult mice fibroblasts^7^. In humans, homologous factors could induce neural conversion in fibroblasts at a very low efficiency^8^, which could be improved by the addition of other factors such as Ngn2, Sox2, NeuroD1/2, miR-9/9*, and/or miR-124^6,9,10^. The efficiency of some of these methods has been increased by concomitant knockdown (KD) of the REST complex^6,10,11^. Another approach involves knockdown of PTB^12,13^. Other factors have been used to convert to specific neural subtypes^9^.

The experimental successes in the field of neural reprogramming indicate that, although the specialized identity of the cell constitutes many different aspects of cellular form and function, it is fundamentally controlled by only a few key regulators. Much interest has gone to identifying these key regulators and their roles, but the attempts to integrate this knowledge into a holistic understanding of the gene interaction network are still in the early stages. A quantitative model that captures the essential properties of the neural conversion mechanism represents an invaluable tool for suggesting new experiments and more efficient reprogramming methods. Researchers have engaged in such modelling in closely related fields such as the study of pluripotent cell commitment and reprogramming somatic cells to pluripotency. The resulting models successfully compress many experimental results into a single framework that can be interpreted by researchers, provides further experimental predictions, and can be used to suggest new experiments^12,13,14^. With the goal of understanding direct neural reprogramming, an initial network hypothesis has been published^13^, but its ability to reproduce and explain the behaviour of the cell has not yet been explored in a quantitative way.

In this study, we built a quantitative model of the gene regulatory system governing direct reprogramming to neurons, based on known gene interactions that have been described in literature. We then measured the expression levels of key transcription factors at different time points during a neuronal reprogramming experiment, and used the resulting experimental data to evaluate the literature-based model. We found that this literature-based model was not able to explain the experimentally observed dynamic behaviour. However, adjustments to the network topology could drastically increase the match between model simulations and experimental observations. In particular, all the adjustments that were shown to substantially improve the model were characterized by an activation from nPTB to PTB, not previously described in literature. Further, we showed that this novel interaction allows the gene regulatory network to successfully predict the reactions of the cells to various overexpression and knockdown perturbation experiments, including several different neuronal conversion strategies. In contrast, models based on network hypotheses without this crucial interaction could not reproduce the correct system behaviour. Together, these results suggest that nPTB plays an activating role in the expression of PTB, for example by blocking the negative self-regulation of PTB. In addition, we present a novel approach to dissect the model to reveal the influence of individual interactions during the conversion process, potentially illuminating internal mechanisms of the network that are hard to observe in the lab. By incorporating the mechanisms of different conversion methods into one explanatory framework, this work aims towards an integrated and predictive understanding of cellular decision making in direct neuronal reprogramming.

## Results

### Measured transcription levels during neuronal reprogramming

We have previously shown that cells that are converted by knocking down REST adopt a transcriptome that more closely resembles that of a neuron, compared to the reprogramming approach involving the overexpression of miR-9/9* and miR-124^6^. However, the combination of both the REST knock down and miR-9/9* and miR-124 overexpression promotes neuronal maturation^10^. By building on this work, we sought to have a better understanding of how the different components of the gene regulatory network (GRN) controlling direct conversion to neurons interact. We thus generated induced neurons (iNs) through the knockdown of REST, followed by the forced expression of the transcription factors Ascl1, Brn2 (Figure 1a). This reprogramming protocol, which leads to a striking change towards a neuronal morphology during the first 25 days of conversion (Shrigley et al., 2018), generates iNs that express synaptic markers such as synapsin, synaptophysin and that are electrophysiologically functional (Drouin-Ouellet, Lau, et al. 2017), and genes of specific neurotransmitter phenotypes, including somatostatin (SSTR1), GABAergic (GABRA1), glutamatergic (GRIA2), acetylcholinergic (CHRMA43), and dopaminergic (DRD1)^10^. The neuronal identity of the reprogrammed cells was also confirmed in this study by the expression of the mature neuronal markers TAU and MAP2 at day 25 (Figure 1b). This reprogramming strategy generates iNs at an efficiency of 43.9 ± 8.9%, as calculated by the percentage of the total number of TAU^+^ cells over the number of starting fibroblasts, as well as a purity of 44.1 ± 1.1%, as calculated by the percentage of the number of TAU^+^ cells over the total number of cells in the dish at day 25 (Figure 1c). We measured mRNA concentrations of known key transcription factors in the fibroblast before conversion, as well as three days following the REST knockdown, and 8 hours, 1, 2, 3, 5, 7, 14, and 21 days following the induction of viral expression vectors (Figure 1d). Endogenous Ascl1 and Brn2 were measured separately from their viral counterparts. No endogenous Brn2 transcription was observed.

**Figure 1:**
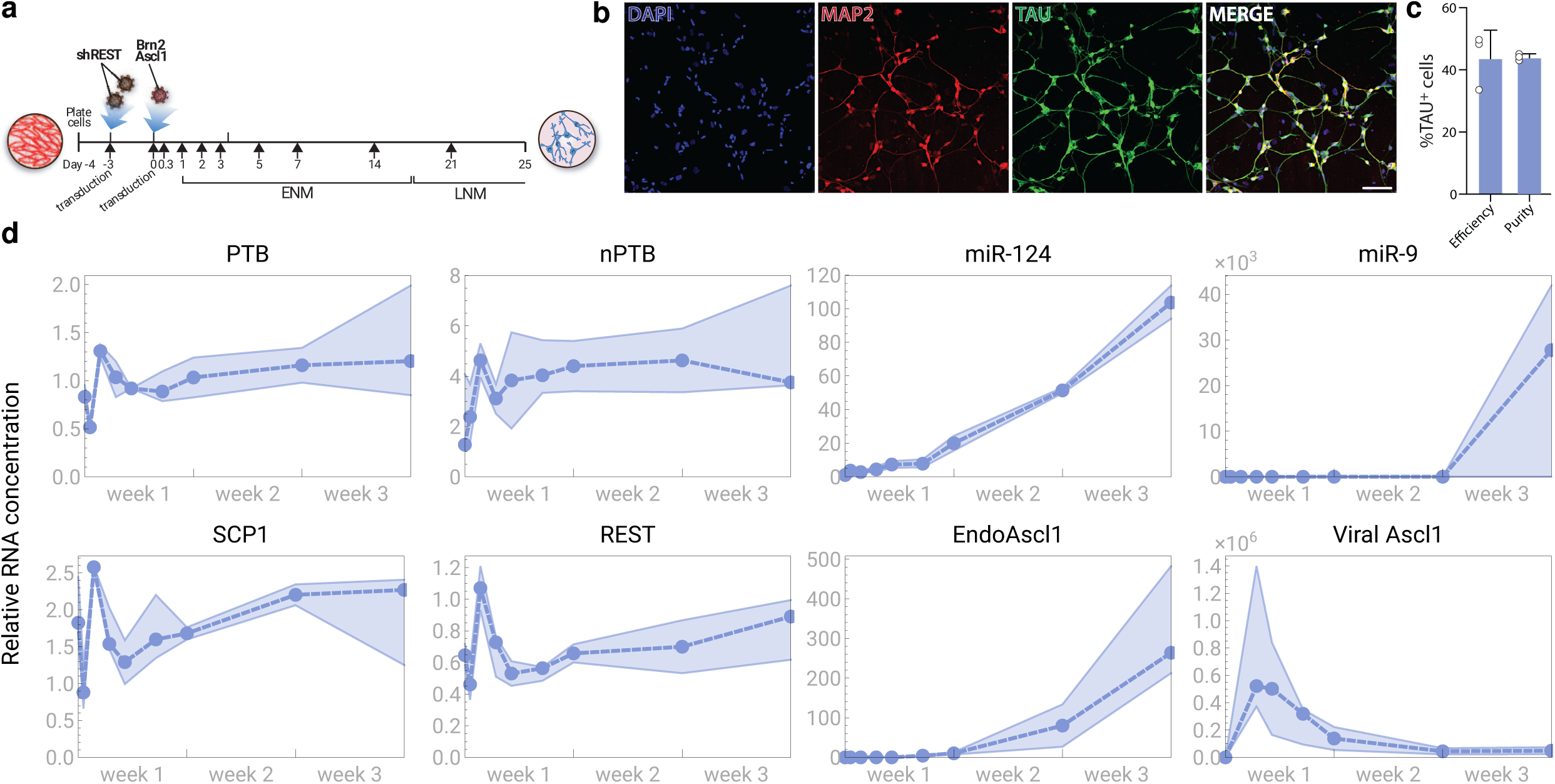
Measured transcription levels during neural reprogramming. (**a**) A schematic representation of the reprogramming process. (**b**) Immunofluorescence staining of MAP2 (in red) and TAU (in green) showing reprogrammed iNs at day 25 post transduction. Cells are counterstained with DAPI (in blue). Scale bar = 100 µm. (**c**) Quantification of TAU+ cells at day 25 post transduction. (**d**) Measured transcription levels. The dashed line represents the median of three replicates, surrounded by a shaded region between the minimum and maximum values at each time point. The measurements are normalized so that a value of 1 is equal to the fibroblast level, or to the detection limit in the case of Ascl1, which was not observed in fibroblasts. Abbreviations: ENM: early neuronal medium; LNM: late neuronal medium

Our measurements showed intense fluctuations in the expression levels during the first five days, followed by stagnation or monotonous increase during the next two weeks. This is consistent with previous observations that commitment to neuronal development occurs quickly in the reprogramming process in human cells, followed by a longer period in which downstream genetic and epigenetic control mechanisms are rewired^15,16,17^. The observed dynamics are typical of a system responding to an initial input shock before settling into a new stable regime.

We noted that the viral expression was not strongly correlated with the other expression curves, and even at lower observed levels was several orders of magnitude above fibroblast concentrations. Throughout the rest of this study, we assumed that the regulatory mechanisms depending on Ascl1 were saturated during the time frame where it was virally expressed, and considered the absence or presence of Ascl1 as an on-off switch in our models.

### Testing the literature-based network

In the last decade, there has been considerable interest in the role of several key factors that play an important role in establishing and maintaining neural identity. We integrated this existing knowledge (Table 1) into a gene regulatory network suitable for comparison with our data set (Figure 2a). This GRN is based especially on previous work by Xue et al.^16,18,19^. We focused here only on neuronal conversion, not maturation.

**Table 1:**
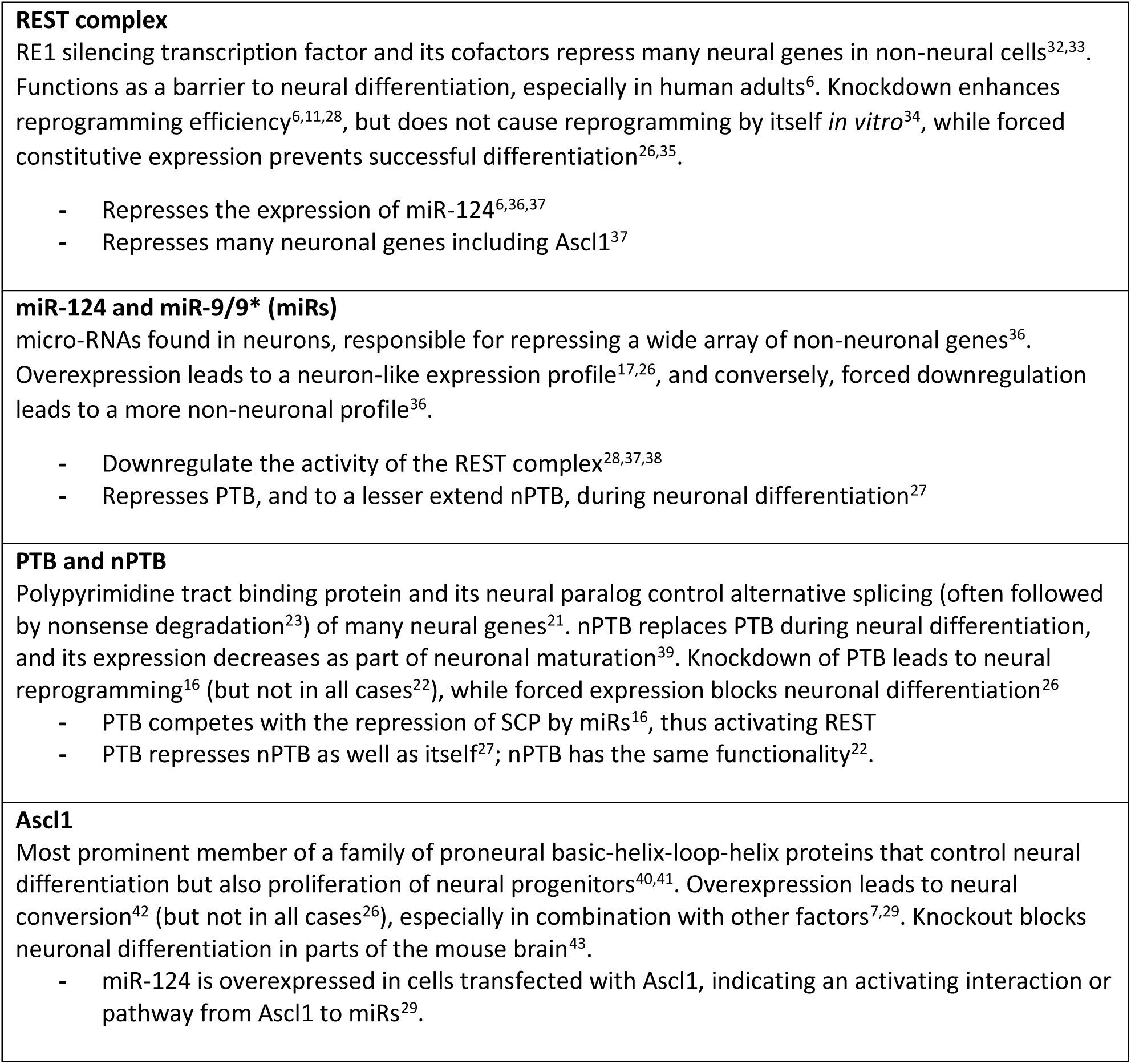
Genes in the core regulatory network. These factors have been indicated as key to establishing and maintaining neural identity.

**Figure 2:**
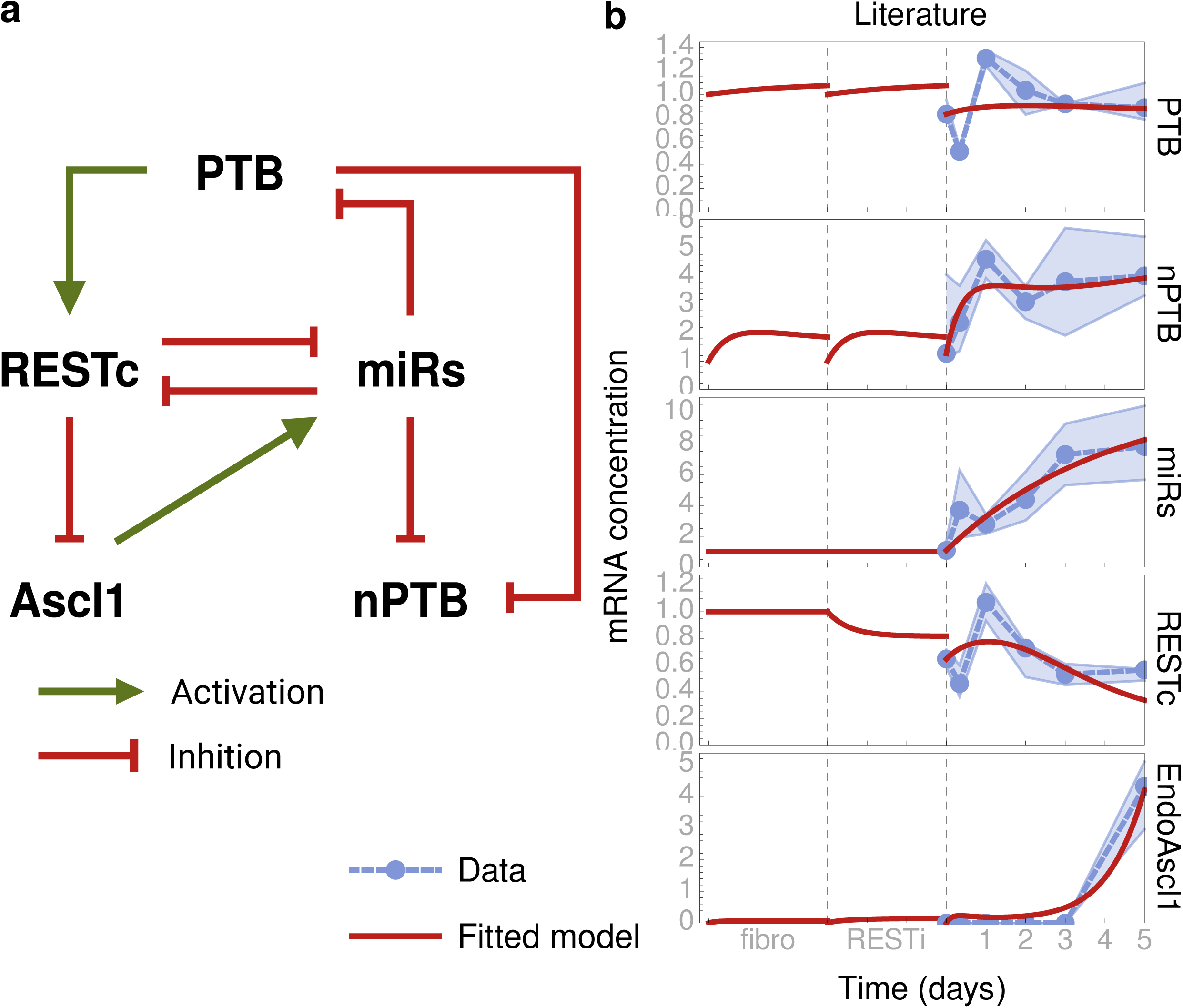
Known interactions are not sufficient to explain observed reprogramming dynamics. (**a**) Literature-based network topology. Green pointed arrows represent activations, red flat arrows represent inhibition interactions. (**b**) Predicted transcription levels according to quantitative model based on gene network in A, fitted to our experimental data. Time scale is divided in three phases, corresponding to fibroblast, REST knockdown, and conversion stages of the experiment.

Two nodes in our network represent the combined expression of multiple physical regulatory elements: the miRs node represents miR-124 and miR-9/9* while the REST complex represents its multiple components, including the REST gene and SCP1 cofactor. This approach was motivated by high similarity between the relevant transcription levels during our experimental time frame (correlation coefficient of median concentration levels *r* = 0.89 (miR-124–miR-9) and *r* = 0.89 (REST–SCP1)), especially during the first five days (*r* = 0.97 (miR-124–miR-9) and *r* = 0.93 (REST–SCP1)). Thus, there was almost no quantitative distinction between these regulatory elements during the conversion process that we observed. Experimental data gathered from other conversion methods could help distinguish the action of these factors in the future.

We developed a quantitative computational model for the GRN in Figure 2a, using the Shea-Ackers formalism^20^, yielding a system of ordinary differential equations (see Methods) that predicted the dynamics of RNA concentrations over time for each node in the network. The interaction strengths and reaction rates were unknown parameters in these equations; we used an evolutionary algorithm to find the parameter values which yielded the closest match between predicted and observed transcription dynamics (see Methods). The model was executed in three modes corresponding to the three stages of our experimental conversion process: the initial fibroblast cell stage, the REST knockdown stage, and the conversion stage with Ascl1 and Brn2 overexpression. The switch between the three modes was modelled by manipulating two parameters representing the action of external factors i.e. REST knockdown and Ascl1 overexpression, respectively. System parameters were optimized to maintain a steady state in the fibroblast stage, and to exhibit dynamics that fit to the observed transcription levels in the other two stages.

When fitting our simulation outcomes to the experimental time series data, we found that our initial model based on interactions described in literature was not able to explain the dynamic behaviour of the system (Figure 2b). Two observations stood out as possible explanations for the inability of the model to fit the data.

First, while the behaviour of the system during the first five days of observation appeared to be dominated by the shock response to the sudden addition of the viral vector-delivered Ascl1 to the system, there was no interaction in the network that could explain these heavy fluctuations in expression levels. In particular, the strongest fluctuations occurred during the first day, when PTB, REST, and miRs drastically switched from increasing to decreasing transcription levels, or vice versa. None of these factors had an activator or inhibitor in this GRN model that reached a low or high enough level to drive these drastic changes in direction. Because nPTB was the only factor affecting the rest of the network whose transcription levels appeared to be changing steadily during this time period, it was inviting to speculate that an unknown interaction from nPTB to another node in the system was responsible for the sudden change in behaviour.

Second, we noted that, in this GRN, Ascl1 was controlled exclusively by inhibition from the REST complex. Therefore, it was activated only by constitutive transcription factors not present in the network. However, at time points where Ascl1 was expressed, we did not observe a uniquely low REST expression (Figure 1c). This suggested that another regulatory process might have been necessary to accurately explain the expression patterns of Ascl1 in the context of reprogramming. This regulatory process could have affected Ascl1 either directly, or indirectly by interfering with other components of the REST complex. The absence of such a regulatory interaction in our model could have driven other interaction strengths to compensate, potentially constraining all system parameters to a poor fit. An alternative explanation for the lack of Ascl1 expression during the first day is that the low REST transcription is not sustained for long enough to sufficiently affect protein concentrations.

We also noted that, in addition to the interactions shown in Figure 2a, previous publications have described negative feedback between PTB and nPTB, as both proteins have been found to inhibit the correct splicing of their own and each other’s mRNA transcripts^21,22,23^. Except for the inhibition of PTB by nPTB, these interactions have not received much attention in literature as causal elements influencing the neural reprogramming process, and they are not represented in our literature-based model.

### Comparison of alternative network topologies

To investigate whether a small modification to the GRN topology could improve the match between predicted and observed transcription levels, we explored several alternative network hypotheses.

Based on the above observations, we initially tested for two additional regulatory control mechanisms if they could improve model performance. First, we tested the hypothesis that the fit is strongly influenced by the constraint that the REST complex is most active near the end of the five days period, which is necessitated by the fact that REST is the only activator of Ascl1. To achieve this, we fitted an alternative model where Ascl1 was additionally activated by miRs. Because miRs are also at their highest level at the end of the five days, this activation (with appropriate parameters) is sufficient to fit Ascl1, without putting hard constraints on REST. As a second model alternative, we introduced inhibitions in the PTB/nPTB switch (PTB ⊣ PTB, nPTB ⊣ nPTB, nPTB ⊣ PTB; the interaction PTB ⊣ nPTB is present in all models). As shown in Figure 3a top, the Ascl1 activation resulted in a small improvement to the model fit, while adding the negative feedback between nPTB/PTB had no effect.

**Figure 3:**
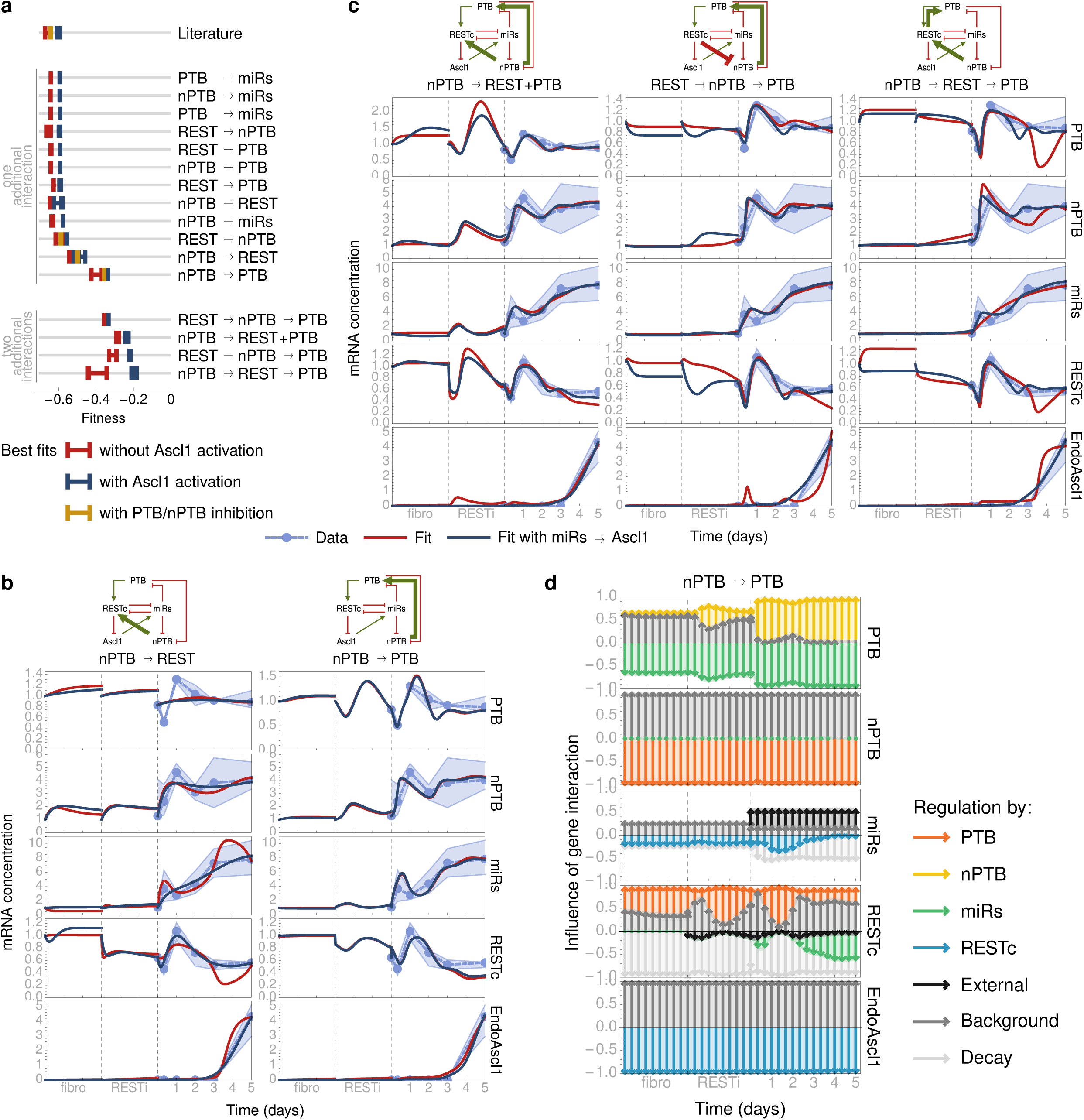
Model performance increases drastically when nPTB activates PTB. (**a**) Model performance as measured by the objective (fitness) function of the optimization procedure, that is, the negative fitting error. Performance of the three best fits is shown. Models are grouped by the hypothetical interaction they have in addition to the literature-based network, where → denotes activation, and ⊣ denotes inhibition. (**b**) The time evolution of the RNA concentrations predicted by the nPTB → REST and nPTB → PTB models. Time scale is divided in the three experimental phases as in Figure 2b. (**c**) The time evolution of the DNA concentrations in the best models with two additional interactions. (**d**) Dissection of the simulated time evolution in the nPTB → PTB model (see (**b**)), decomposing the transcription rates of each factor into the separate contributions of its activators and inhibitors. Activating interactions are represented by arrows pointing upwards from the zero line, inhibiting interactions point downwards. Arrow length indicates relative influence of the regulator. See Methods for an extended description. nPTB and Ascl1 are controlled by changes in inhibition regulation that are too small to see on the figure.

To further explore whether the fit could be improved by a small modification to the system, we investigated alternative models where single interactions were added between the four factors PTB, nPTB, miRs, and the REST complex. We excluded models with self-interactions (interactions from a node to itself) and double interactions (two interactions with the same origin and target). Results are shown in Figure 3a one additional interaction (middle) part. For each of these models, we also tested whether an additional Ascl1 activation could improve fit, confirming that the improvement conferred by this modification is small. The nPTB/PTB switch self-inhibition, which did not improve the literature model, was also tested for the three best models, also confirming that these inhibitions do not improve model performance despite adding several degrees of freedom.

Most of these models with one additional interaction did not provide a valuable improvement compared to the literature-based network. In other words, adding more degrees of freedom to the model selection did not affect the ability to fit transcription data in most cases, making the few exceptions notable. Among all the tested models, by far the lowest fitting error was found when nPTB activated PTB (nPTB → PTB). To a lesser extent, fitting also improved when nPTB activated REST (nPTB → REST). The expression levels predicted by these models over the course of the conversion experiment are shown in Figure 3b. These outcomes corroborated the hypothesis that the accumulation of nPTB during the first day after introduction of virally delivered Ascl1 activates an undiscovered interaction that is responsible for much of the observed system dynamics. Note that the nPTB → PTB and nPTB → REST models are coherent in the sense that each of these interactions indirectly implied the other through an intermediate reaction chain: nPTB → [PTB →] REST and nPTB → [REST ⊣ miRs ⊣] PTB, where the number of inhibition interactions in the chain is even in both cases.

We next tested if the fitting error could be further improved by adding two hypothetical interactions to the network (not including the additional activation of Ascl1 by miRs)), shown in Figure 3a (bottom part) and Figure 3c. As before, there were only a few cases where adding more hypothetical interactions improved upon simpler models. The fitting error could be reduced by combining the two best single interactions (nPTB → REST+PTB), and by combining the activation from nPTB to REST with an activation from REST to PTB (nPTB → REST → PTB), so that the nPTB → PTB interaction is again included implicitly. We also saw an equally large improvement when REST inhibited nPTB in addition to nPTB activating PTB (REST ⊣ nPTB → PTB). Notably, this last model could be further supported by the presence of an RE1 binding site close to the nPTB locus^24^. These models have in common an additional interaction between the PTB/nPTB switch and the REST complex, which is also present in the second most performant model with only one hypothetical interaction (nPTB → REST). Thus, such an interaction is highly compatible with our observations. Fitted parameter values for this and other models are shown in Supplementary Table 2.

Overall, improvement to the fit between simulation outcomes and experimental data was by far most pronounced when a hypothetical activation from nPTB to PTB was introduced to the network. The specificity of this improvement suggests the possibility that such an interaction contributes to the dynamics of neuronal reprogramming using the strategy of Ascl1 overexpression coupled with REST knockdown.

The equations in these quantitative models were built up from the interactions in a gene regulatory network using a systematic procedure. As a result, different parts of the equations are identified with specific gene interactions. By comparing the magnitude of partial expressions that represent contributions of individual gene interaction, we can measure and visualize the relative influence of each activator or inhibitor on the total transcription rate of each gene at different time points during a simulation. This dissection of the model allows us to identify crucial gene interactions by directly observing the (model) system’s internal mechanisms in full detail. In contrast, in the lab it is only possible to observe the behaviour of the network as a whole, for example in response to overexpression or knockdown of genes, leaving the contributions of individual interactions to inexact inference. Figure 3d shows such a dissection analysis for a simulation of the nPTB → PTB model (the corresponding timed expression profile shown in Figure 3b). See Methods for the analytical details of this analysis. The analysis shows, for instance, that the training data taught the model that the inhibition from miRs has a negligible impact on nPTB expression. This interaction may still contribute to the behaviour of the real cell, but apparently its function is redundant (i.e. is functionally equivalent to the inhibition by PTB) in the conditions of our conversion strategy.

### Model performance in overexpression and knockdown scenarios

Having confirmed that several network hypotheses are highly compatible with gene expression data during one conversion process, we set out to investigate whether the fitted models would also reproduce the behaviour of the cell in response to other stimuli. To this end, each model was subjected to simulated overexpression (OX) or knockdown (KD) of factors that are known to cause or block neural conversion (see Table 1). The results could then be compared to experimental observations. Since these qualitative data are independent of our training data, they are suitable for model testing and validating comparison.

In order to simulate analogs to these experiments in our models, we explored the effect of modifying relevant parameters on the system behaviour. Viral induction of Ascl1 and REST knockdown were explicitly represented in the system equations and so could be controlled by manipulating the dedicated system parameters. This is the same method used when switching between simulating the three stages of the experimental procedure. All other cases of overexpression and knockdown were simulated by increasing (×5) or decreasing (×1/5) the transcription rate coefficient of the factor under consideration. The resulting system equations were then integrated to yield the evolution of transcription levels over time, starting from the fibroblast mRNA expression level as an initial state. Because many direct reprogramming methods operate at low efficiency, the deterministic integration representing bulk average expression levels of a population was supplemented by a stochastic approach, to capture noise in individual cells. We used the Gillespie algorithm to produce stochastic simulations over a simulated time span of 14 days^25^. Endogenous expression of Ascl1 was taken as a convenient indication that a simulated cell had committed to a neuronal fate; a maximum endogenous Ascl1 value of 1 is taken as the threshold value to indicate conversion. Results are shown in Table 2.

**Table 2:**
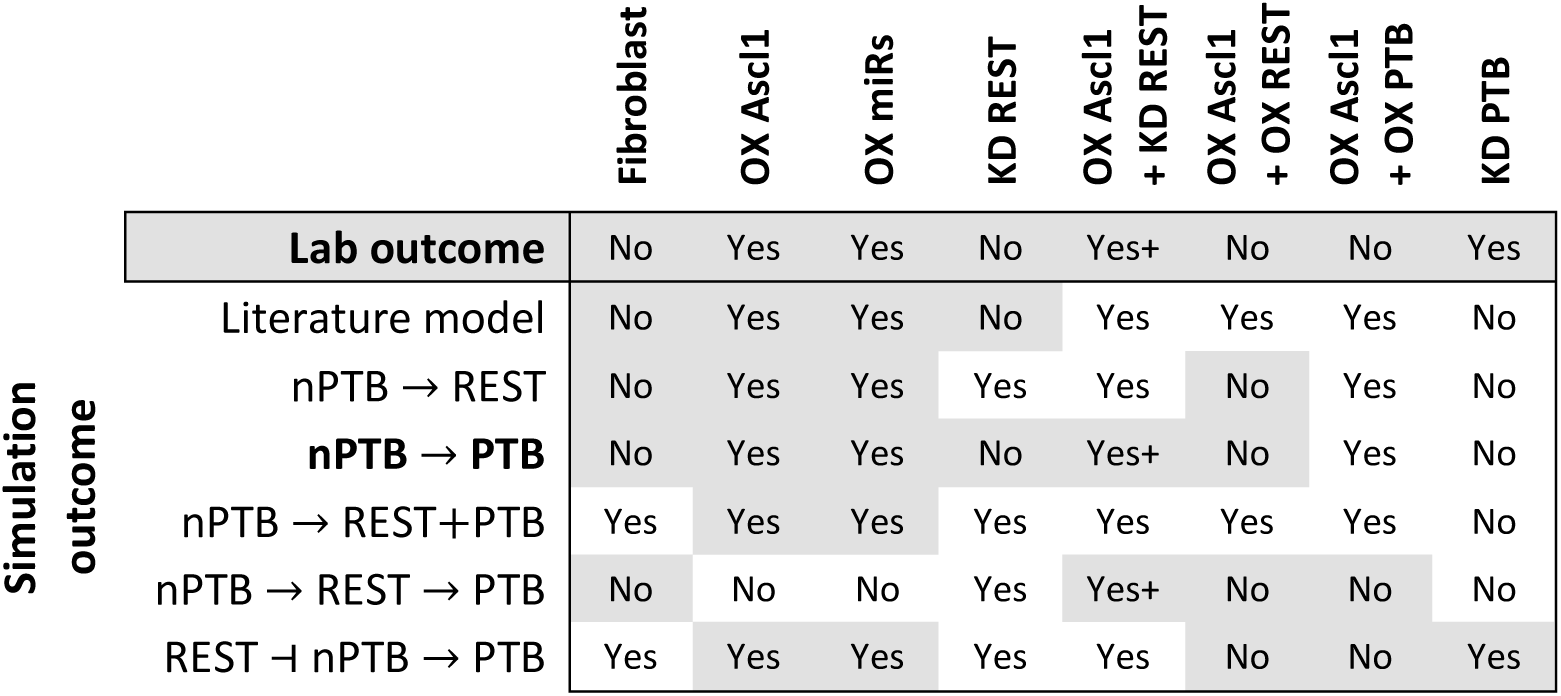
The nPTB → PTB model correctly predicts conversion outcome of most overexpression and knockdown scenarios. Conversion outcome is shown for several OX and KD experiments described in literature (discussed in Table 1), and compared to outcome predicted by quantitative models. Simulated outcome was considered to predict conversion (“Yes”) if Ascl1 expression exceeded 1 at any time point during at least 1 out of 50 simulations, and no conversion (“No”) otherwise. In the case of OX Ascl1 + KD REST, all models predict conversion because it is an explicit feature of the training data. For this reason, a higher conversion efficiency was required (indicated by a plus sign) in this experiment as compared to OX Ascl1. Shaded cells indicate outcomes that correspond to lab experiments. Full simulation results are shown in Supplementary Figure 1.

Out of all the models presented here, nPTB → PTB (Figure 4a) provided the most accurate predictions of system behaviour in different OX/KD scenarios, noticeably improving upon the purely literature-based model. In particular, this model successfully captures the role of REST in the system (see Table 1), showing an increase in conversion efficiency when REST is knocked down, which we have previously validated experimentally^6^, as well as blocking the conversion process when REST is overexpressed during conversion with Ascl1 overexpression (Table 2). The stochastic and deterministic simulation results corresponding to OX/KD scenarios with the nPTB → PTB model are shown in Figure 4b, c.

**Figure 4:**
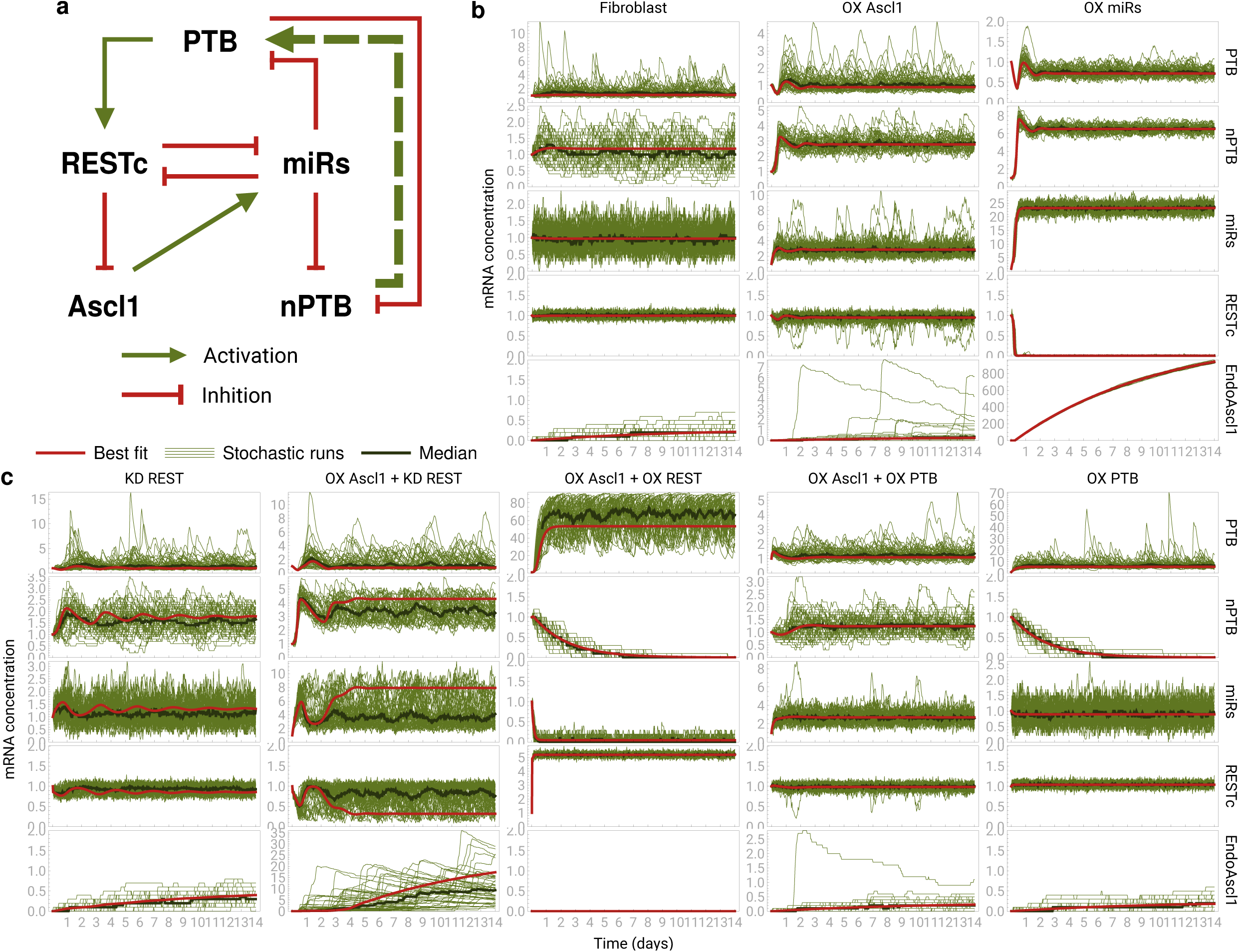
The nPTB → PTB model correctly predicts conversion outcome of most overexpression and knockdown experiments. **(a**) Network topology of the nPTB → PTB model. **(b**,**c)** The deterministic (red) and stochastic (green) simulation outcomes are shown for OX/KD scenarios in the nPTB → PTB model. Similar simulations for all models and experiments are summarized in Table 2 and are shown in Supplementary Figure 1.

### Dissecting the model response to PTB knockdown

Generally, there was a remarkable agreement between the system behaviour predicted by the nPTB → PTB model and experimental observations, despite the fact that the training data cover only a narrow set of circumstances compared to the many perturbation experiments used as validation. The fact that a computational model could learn the outcome of one experiment from training data based exclusively on another experiment, suggests that they operate in the same region of the system phase space, and that these different conversion strategies invoke a similar response from the gene regulatory network. Notably, the exception to this rule is the system response to PTB knockdown (which can cause neuronal differentiation^16^) or overexpression (which is known to block neuronal differentiation with Ascl1 overexpression^26^). This suggests that neuronal conversion induced by PTB knockdown is affected by a different cascade of gene interactions compared to the other conversion strategies. Based on the gene interaction network, one way in which the PTB knockdown strategy stands out is that it affects REST (and through it the rest of the network) by removing a crucial activator (PTB), whereas other conversion strategies directly or indirectly induce miR-124-9/9*, a repressor of the REST complex.

In the nPTB → PTB model, decreasing the PTB transcription rate coefficient (simulating PTB knockdown) resulted in an increase of nPTB without affecting the other factors Figure 5a, left. While different from the expectation that PTB knockdown results in neuronal conversion, this result is corroborated by an earlier report^22^. Dissecting the model showed that, even though REST was no longer activated by PTB, the trained model predicted that constitutive activation of REST expression was enough to maintain nominal concentrations, thus not affecting the rest of the system (Figure 5a and b, left). Interestingly, when the constitutive activation of REST was reduced, the system became permissible to neuronal conversion by PTB knockdown (Figure 5a and b, middle). As a negative control, we confirmed that when constitutive REST activation is decreased without accompanying PTB knockdown, the system is not substantially affected, sustaining a steady state near the expected fibroblast-like transcription levels (Figure 5a and b, right). Thus, it is compatible with our model that some cells are more susceptible to neuronal conversion through PTB knockdown than others, due to variation in constitutive REST expression, meaning expression of components of the REST complex due to factors that were not explicitly discussed in this work.

**Figure 5:**
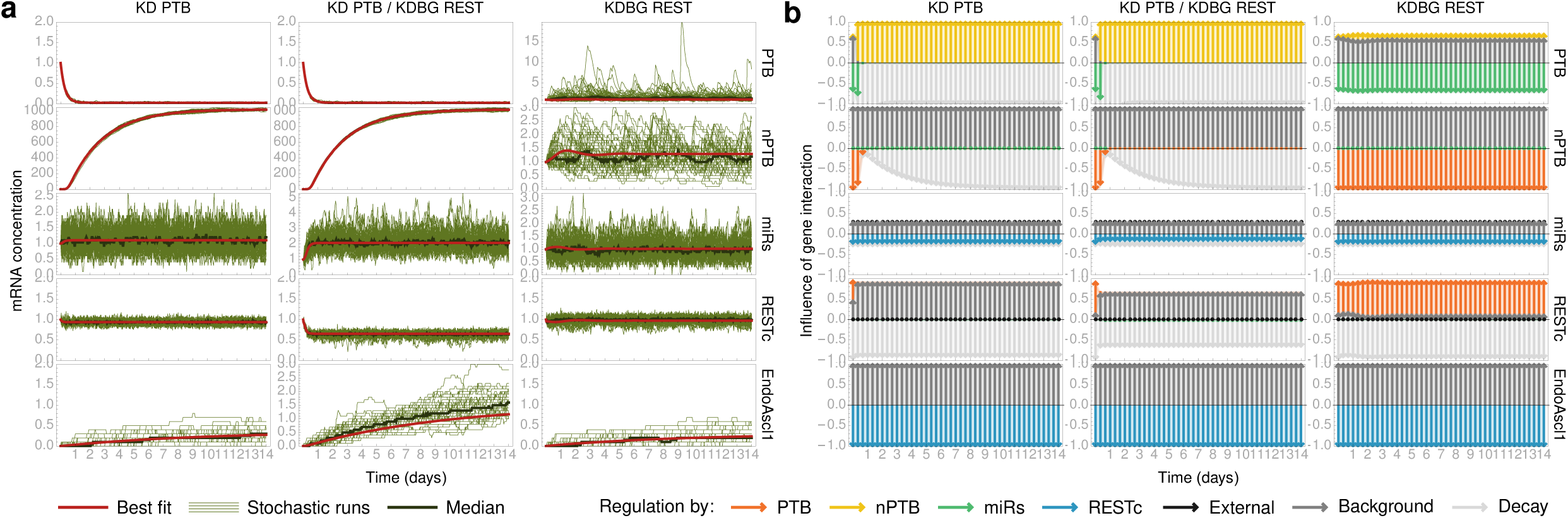
Strong constitutive activation of REST expression blocks neuronal reprogramming through PTB knockdown in models. (**a**) Stochastic (green) and deterministic (red) time evolution of the system. Left: simulated knockdown of PTB. The system remains in non-neural state, as seen by the lack of Ascl1 expression. Middle: combining PTB knockdown with a diminished constitutive REST activation induces Ascl1 expression. Right: with regular PTB expression, reducing constitutive REST activation results in a stable non-neuronal profile, with a steady state that strongly resembles base fibroblast levels. (**b**) The influence of different regulatory interactions on the expression of each factor during the deterministic experiments shown in panel **a** (see also Figure 3**d**). See Methods for an extended description.

## Discussion

In this study, we put forward a quantitative model of the core gene regulatory network governing direct reprogramming of human adult fibroblast cells to neurons. Our analysis showed that the known interactions between key transcription factors were unable to explain the full dynamic behaviour that we observed in our experimental data. However, adding a hypothetical interaction to the network where nPTB activates PTB drastically increased the match between measured mRNA expression levels and those predicted by the model. In addition, in contrast to various alternative hypotheses, this model correctly predicted the response of the cell in a range of different overexpression and knockdown scenarios. Taken together, these findings suggest that such an interaction between these two genes may play a role in neuronal conversion. However, the possibilities for how this interaction is manifested biochemically should be interpreted broadly. The model is agnostic of specific regulatory mechanisms, so the cause of the predicted interaction is not necessarily restricted to the transcriptional level and may include mediation by one or more intermediate regulators.

Interestingly, there is a possible mechanism by which nPTB may function as an activator of PTB, in line with known features of PTB and nPTB regulation. Both PTB and nPTB are known to regulate the splicing of PTB mRNA, effectively inhibiting its transcription by inducing nonsense-mediated decay^21,22,23,27^. We have shown that adding these interactions to the network in the form of inhibitions did not improve model performance. However, several publications have noted that nPTB is a weaker splicing regulator than PTB, and that this difference in regulating activity is due to differential recruitment of cofactors rather than RNA binding activity^21,22,27^. By entering in binding competition with PTB to control the splicing of the PTB mRNA, nPTB could effectively weaken the self-inhibition of PTB, thus acting as a net activator. Due to the simplifying assumptions of our mathematical model, such an interaction can only be represented as a direct activation from PTB to nPTB, which may explain our findings. This effect of mutual self-regulation of PTB and nPTB has not previously been explored as a possible causal element in cellular decision making during direct neural reprogramming.

When investigating the response of the different models to simulated overexpression and knockdown experiments, we found a remarkable ability to predict experimental outcomes. The fact that information gained from only one experiment generalizes to other conversion strategies suggests that the different methods operate via the same interaction mechanisms and guide the cell to similar trajectories in the phase space of the system. One consequence of this insight is that we may expect greater conversion efficiencies when compatible strategies are combined. Indeed, several publications have combined REST knockdown, Ascl1 overexpression and miR-124-9/9* overexpression, each of which can induce conversion on its own, with the effect of increasing conversion efficiency and neuronal maturation^6,10,11,28,29^.

Our analysis did not predict that the conversion strategy based on PTB knockdown would result in cells with a neuron-like identity, but instead showed an increase in nPTB expression without affecting other genes. Interestingly, although PTB knockdown has been used successfully in a number of conversion experiments^16,30^, this negative experimental outcome has also been observed in the lab^22^. We provide a speculative but plausible explanation for this variation in results by observing that, in our best model, the conversion was blocked by constitutive activation of REST. According to the model, cells with a lower constitutive REST activation would have similar REST expression levels, but would be more susceptible to conversion after PTB depletion. However, further research is needed to understand the dynamics of gene expression governing this conversion strategy. In particular, gene expression data gathered from cells treated with PTB knockdown could help shed more light on this process.

Overall, this work illustrates that the current understanding of the genes and interactions governing neural reprogramming can be unified into a computational framework that explains much of the observed behaviour of the cell. Our analysis brings a new perspective to some aspects of the cellular decision process that establishes neuronal fate, and demonstrates a novel analytical tool to dissect internal model dynamics. In the future, more detailed quantitative models informed by data from varied experiments can increase the power of this approach, helping to provide a better understanding of the existing conversion strategies, and suggesting improvements that could result in higher yield or faster conversion to neurons.

## Methods

### Experimental Data

#### Cell culture and cell lines

Adult dermal fibroblasts of a 67-year-old healthy female donor were obtained with written, informed consent from the Parkinson’s Disease Research clinic at the John van Geest Centre for Brain Repair (Cambridge, UK). All experimental protocols were approved by the East of Cambridgeshire Research Ethics Committee (REC 09/H0311/88) as well as the University of Montreal *Comité d’éthique de la recherche en sciences et en santé* (CERSES-18-004-D). All experiments were performed in accordance with the Declaration of Helsinki.

For details on the skin biopsy sampling method, please refer to^6^. Primary fibroblasts were expanded and cultured at 37°C in 5% CO_2_ in fibroblast medium (Dulbecco’s Modified Eagle Medium (DMEM) + Glutamax (Gibco) with 100 mg/mL penicillin/streptomycin (Sigma), and 10% FBS (Biosera)). The cells were then dissociated with 0.05% trypsin, spun, and frozen in 50/50 DMEM/FBS with 10% DMSO (Sigma).

#### Viral Vectors and Virus Transduction

DNA plasmids expressing mouse open reading frames (ORFs) for Ascl1 and Brn2 on the same construct (pB.pA, see [6]) as well Lmx1a and FoxA2 [8] in a third-generation lentiviral vector containing a non-regulated ubiquitous phosphoglycerate kinase (PGK) promoter were used (Figure 1a). The knockdown of REST was done using short hairpin RNAs^6,31^. All the constructs have been verified by sequencing. Lentiviral vectors were produced and titrated as previously described^6^. Transduction was performed at a MOI of 10 for the pB.pA vector and an MOI of 5 for all of the other vectors.

### Neural reprogramming

For direct neural reprogramming, fibroblasts were plated at a density of 26 300 cells per cm^2^ in 24-well plates (Nunc) coated with 0.1% gelatin (Sigma). On the next day, cells were transduced with shRNAs against REST and the medium was changed the following day. Three days after this viral transduction, cells were transduced with the pB.pA, Lmx1a and FoxA2 constructs. Fibroblast medium was replaced by neural differentiation medium (NDiff227; Takara-Clontech) supplemented with growth factors at the following concentrations: LM-22A4 (2 µM, R&D Systems), GDNF (2 ng/mL, R&D Systems), NT3 (10 ng/µL, R&D Systems) and the small molecules CHIR99021 (2 µM, Axon), SB-431542 (10 µM, Axon), noggin (0.5 µg/ml, R&D Systems), LDN-193189 (0.5 µM, Axon), as well as valproic acid sodium salt (VPA; 1mM, Merck Millipore) and db-cAMP (0.5 mM, Sigma), three days after the second transduction. Half of the neuronal conversion medium was replaced every 2-3 days. 18 days post-transduction, the small molecules were stopped, and the neuronal medium was supplemented with only the factors LM-22A4, GDNF, NT3 and db-cAMP until the end of the experiment.

#### qRT-PCR analysis

Total RNA, including miRNA, was extracted from cells from the same line at different stages of conversion (Figure 1a) using the micro miRNeasy kit (Qiagen) followed by Universal cDNA synthesis kit (Fermentas, for RNA analysis; Exiqon for miRNA expression). Three reference genes were used for each qPCR analysis (ACTB, GAPDH and YWHAZ). Primer sequences can be found in Supplementary Table 1. LNA-PCR primer sets, specific for hsa-miR-9-5p, hsa-miR-124-3p and hsa-miR-103 (the latter used as normalization miRNA), were purchased from Exiqon and used for the miRNA qPCR analysis. All primers were used together with LightCycler 480 SYBR Green I Master (Roche). Standard procedures of qRT-PCR were used, and data were quantified using the ΔΔCt-method. Analyses were performed in triplicates at each time point.

#### Immunocytochemistry, quantification, and imaging

Cells were fixed on day 25 in 4% paraformaldehyde, permeabilized with 0.1% Triton-X-100 in 0.1 M PBS for 10 min. Thereafter, cells were blocked for 30 min in a solution containing 5% normal serum in 0.1 M PBS. The following primary antibodies were diluted in the blocking solution and applied overnight at 4°C: mouse anti-TAU clone HT7 (1:500, Thermo Scientific), chicken anti-MAP2 (1:10,000, Abcam). Fluorophore-conjugated secondary antibody (Jackson ImmunoResearch Laboratories) was diluted in blocking solution and applied for 2 hrs. Cells were counterstained with DAPI for 15 min followed by three washes in PBS. The total number of DAPI^+^, TAU^+^ and MAP2^+^ cells per well were quantified and imaged using the Cellomics Array Scan (Array Scan VTI, Thermo Fischer), which is an automated process ensuring unbiased measurements between groups. Applying the program “Target Activation”, 289 fields (10 X magnification) were acquired in a spiral fashion starting from the center.

#### Building quantitative models from gene networks

The Shea-Ackers formalism is a mathematical model that describes the activity of a regulatory sequence as a function of the concentrations of activating and inhibiting factors^20^. Assuming that the regulated gene is active when the binding sequence is occupied by an activator (∈ *act*), and inactive when it is unoccupied or in the presence of an inhibitor (∈ *inh*), the formalism borrows techniques from statistical mechanics to compute the average activity at the site assuming thermodynamic equilibrium:

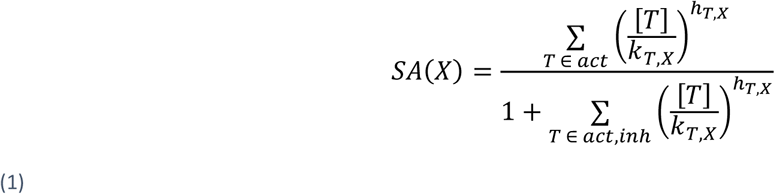

where *k*_*T,X*_ are dissociation constants describing the interaction strength between factor *T* and the regulatory sequence of *X*, and *h*_*X,T*_ is a non-linearity parameter representing cooperativity between multiple factors of the same type.

The genes in our model may be affected by other regulatory elements besides the factors in our GRN model, such as constitutive transcription factors. The concentration of these background factors is taken to be constant. Through a change of variables, extraneous constant terms in the numerator and denominator of the Shea-Ackers fraction representing these background activators and inhibitors can be replaced by a single parameter *b*_*X*_ added to the numerator and denominator (thus acting as an activator). If *SA*(*X*) is multiplied by a free variable, as we do in the transcription rate equations below, the background parameter *b*_*X*_ is also redundant for genes that are not activated by other non-constant elements of the model. In such cases, the influence of background factors on *SA*(*X*) is simply that the otherwise empty sum in the numerator is replaced by 1.

For REST and miRs, formula (1) must also take into account activation and inhibition due to the externally added factors shREST and viral Ascl1, respectively. Because the effects of these external factors are considered either zero or constant, depending on the stage of the experiment, redundant parameters can again be simplified to the constants 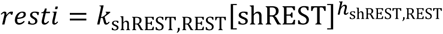 and 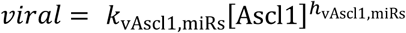, appropriately multiplied by 0 or 1. Viral Ascl1 is modelled as constant with respect to time, because the measured values are at least 10^5^ times greater than the baseline value for fibroblasts during the time frame under consideration, and the system behaviour shown in Figure 1c does not suggest a fluctuating regulatory influence of Ascl1.

Under the assumption that the transcription rate is proportional to the activity of the regulatory site (defined by equation (1) as the average fraction of time the site is occupied by an activator), and that each gene product undergoes exponential decay, the change in gene product concentration over time can be described as

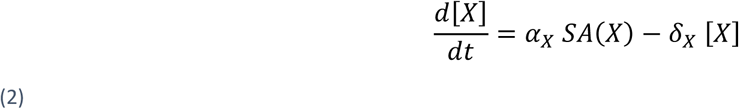

where *α*_*X*_ and *δ*_*X*_ are scaling coefficients for the rate of transcription and decay, respectively (decay half-life is ln(2)/*δ*_*X*_). Using one such equation for each factor in the network yields a system of interacting differential equations that can be numerically integrated to yield a time evolution of concentration levels 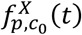 for each factor *X*, given initial concentration values 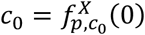 and an appropriate parameter set *p* = {*α*_*X*_, *δ*_*X*_, *b*_*X*_}_*X*_ ⋃ {*k*_*T,X*_, *h*_*T,X*_}_*T,X*_ ⋃ {*resti, viral*}. The full ODE system for the literature-based network model is given in equations (3).

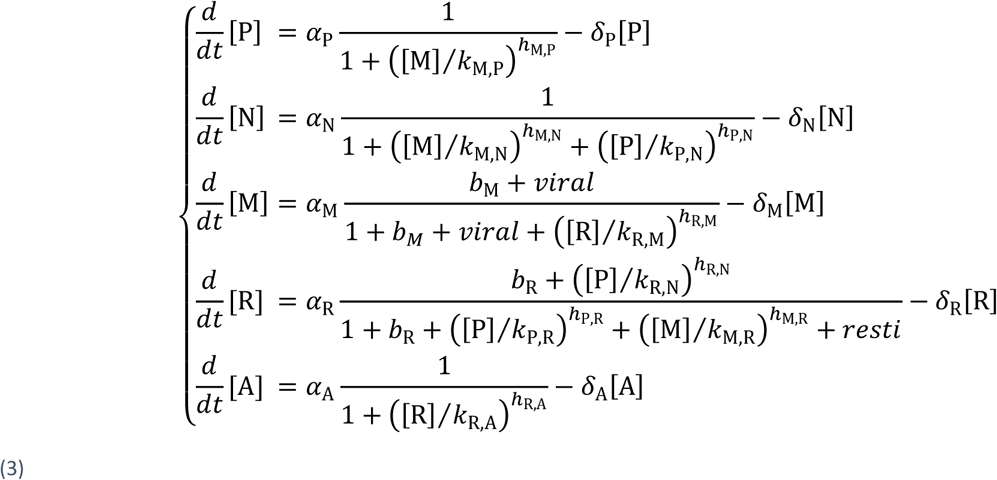

The Shea-Ackers formalism is derived with the assumption that all activations and inhibitions are physically realised by binding competition on a gene’s regulatory sequence. Moreover, the rate equations above assume that there is transcription at a constant rate whenever an activator is bound, thus ignoring differences in the rate at which transcription factors recruit the transcription machinery. Overlooking the complexity of different regulatory mechanisms allows for a mathematical approximation that can describe a wide range of dynamic behaviours that are realistic for different gene interaction mechanisms while using only a small number of parameters.

#### Parameter fitting

A gene regulatory network model produces a good match for the data if there is some parameter set *p* that leads to predicted expression curves 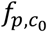 that closely match observations. Finding such a parameter set is an optimization problem. We obtained the best results using a real-valued genetic algorithm, evolving a vector of real numbers which each represent a system parameter. The algorithm uses a population size of 100, uniform cross-over, and a mutation operator that modifies a small number of genes by multiplying with a log-normally distributed random variable. Specifically, if *m*(*p, σ*) represents a random distribution that gives a probability (1 − *p*) of being 1 and a probability *p* of being distributed as exp(𝒩(0, *σ*)), then mutation was implemented by multiplying each variable with independent random values from the distribution *m*(2/*n*, 1.01) * *m*(2/3*n*, 1.05) * *m*(1/5*n*, 1.5), where *n* is the number of genes. The initial population is composed of individuals with genes chosen uniformly from the unit interval, but more complex models were also run using parameters from simpler models as initial values. Each real-valued gene represents a system parameter. Hill parameters *h*_*T,X*_ are constrained to a maximum value of 4 by transforming the internal representation 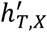 with the function 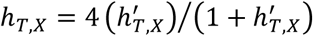 before obtaining the value used in the system equations. The other parameters are used directly as they are in the genome. This means they are unbounded (though the mutations will never yield negative values).

The fitness (target) function of the optimization procedure is based on a standard sum of squared errors (SSE) with smoothness regularisation. For a given set of parameters, the system of differential equations is numerically solved in three stages matching our experimental setup: a fibroblast-like stage *f*_*p*,fibro_ where *resti* and *viral* are set to 0 and initialised with concentration levels observed in the fibroblast; a REST inhibition stage *f*_*p*,RESTi_ where *resti* is included and also initialised with fibroblast concentrations; and a conversion stage *f*_*p*,conv_ initialised with concentration levels observed just before introduction of the viral vectors, where both *resti* and *viral* are used.

The SSE term measures the difference between the system solution *f*_*p*,conv_ and our observed data points during the first five days of conversion termed X in the following equation. The data values used here are the median values of the triplicate observations in each of the six time points. The first data point at *t* = 0 is compared to the final value 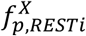 (3 days) of the REST inhibition stage, while the other time points are compared to the conversion stage 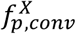.

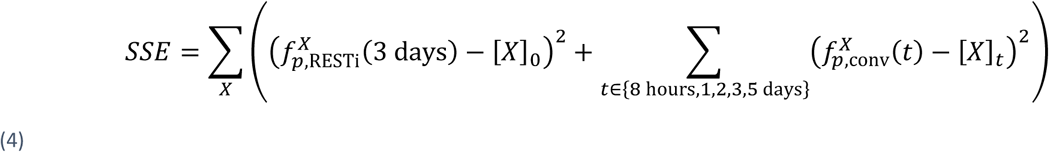

A pure SSE approach often resulted in highly oscillating fits, which closely matched the data but fluctuated erratically in between. To guide the parameter search towards more realistic solutions, we added a smoothness regularisation term of the form:

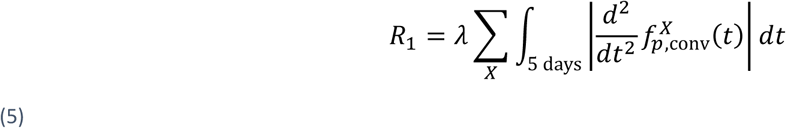

Finally, an important property of the system being modelled is that fibroblasts do not spontaneously convert to neurons, i.e. the mRNA concentrations should remain stable when initialized near the observed fibroblast baseline and when both *resti* and *viral* are set to zero. This is enforced with another term in the fitness function, punishing any changes in concentration during a three-day simulation of the fibro stage.

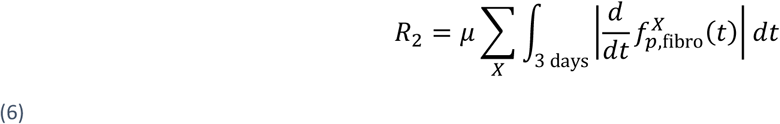

We obtained best results with 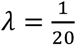 and 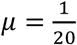. In the evolutionary algorithm, parameter sets with a higher fitness are considered better, so the fitness is set equal to the negative value −(*SSE* + *R*_1_ + *R*_2_).

Two nodes in our networks represent a combination of more than one gene for which expression was measured. We fitted the miRs network node to transcription data from miR-124, and the REST complex network node to data from the REST gene. We obtained qualitatively identical results when using data from miR-9/9* or SCP1, respectively.

### Stochastic simulation

Stochastic simulations were performed using a Gillespie algorithm^25^. Noise levels were adjusted by modifying the reaction formulas: the stoichiometric coefficients were multiplied by a noise level *η*, and reaction rates correspondingly divided by *η* to maintain the same expected number of molecules added or removed per time unit. Our data is a bulk average and does not suggest a value for *η*. In the figures shown here (Figure 4b, c, Figure 5a), its value was set to 0.1. However, two adjustments were made. First, PTB and REST have a (relative) concentration range close to 1, whereas the other factors have typical concentration levels closer to 5. For this reason, transcription and decay reactions for these two factors were transformed with an *η* that was five times lower. Second, these same factors also tended to fit with much higher reaction rates, i.e. both higher *α* and *δ* in (2). To obtain more similar noise levels, *η* for these reactions was adjusted by an additional factor of 1/5, resulting in a total of *η* = 0.004 for PTB and REST, and *η* = 0.1 for nPTB, miRs and Ascl1.

Overexpression and knockdown were simulated by modifying the appropriate transcription rate coefficient. Specifically, for the overexpression, resp. knockdown, of a factor *X*, the corresponding transcription rate coefficient *α*_*X*_ was multiplied by 5, resp. 1/5. Overexpression of Ascl1 and knockdown of miRs were instead controlled by multiplying the values *viral* and *resti* with 0 or 1, as described above. In the case of PTB knockdown, lack of conversion was also verified when *α*_PTB_ was multiplied by ×1/5000, and at this rate conversion was observed when the background activation of REST was multiplied by 1/5.

### Measuring relative influence of interactions

In order to break down the total transcription rate of each factor into contributions from its activators and inhibitors, the terms in the numerator and denominator of the transcription rate equation (1) were calculated at each time point of a trajectory and plotted, as in Figure 3d and Figure 5b. Activators and inhibitors are represented by arrows pointing upwards and downwards, respectively, with arrow length showing the relative magnitude of that interaction term.

Specifically, the total length of all activation arrows 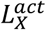 of a gene *X*, at some time point *t*, equals the fraction of the maximum transcription rate that would be realized if the concentration of all inhibitors was zero. The total length of the inhibition arrows 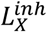 is defined so that the difference between the activation and the inhibition arrows equals the realized transcription activity *SA*(*X*) (equation (1)). This results in the following equations:

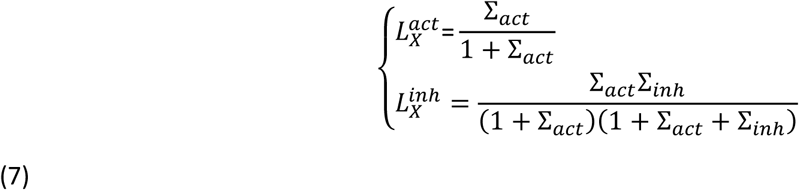

where 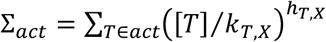 and 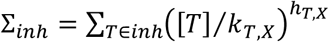 (see equation (1)). Given these total lengths for activator and inhibitor arrows, individual activators and inhibitors *T* are given lengths proportional to their current magnitude 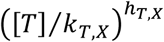, so that the contribution of each inhibitor or activator to the total transcription rate becomes visible.

Finally, an arrow is added to represent the decay term in the equation (2), with a length of 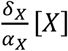 (the decay rate in the same units of *α* as the other arrows). By adding this additional arrow, the difference between the sum of upward and downward pointing arrows shows the rate of change in mRNA concentrations (in units of *α*_*X*_).

## Acknowledgements

JD-O is receiving salary support from the Fonds du Québec en Recherche, Santé (FRQS) and Parkinson Quebec. VO gratefully acknowledges the support of the US National Institutes of Health (USPHS grant R01HL119102). We thank Joachim Eriksson for preliminary work on building the literature-based network.

## Author Contributions

AM, JD-O, VO designed the study. AM, JD-O, VO wrote the manuscript. AM built the model, conducted computational analysis. AM and VD performed parameter optimisation. JD-O produced and analysed the experimental data. RAB provided cells and material. All authors provided inputs and comments on the manuscript.

## Data availability

## Code availability

